# Impact network analysis and the INA R package: Decision support for regional management interventions

**DOI:** 10.1101/2020.11.08.373621

**Authors:** K. A. Garrett

## Abstract

The success of intervention projects in ecological systems depends not only on the quality of management technologies, but also patterns of adoption among land managers. Impact network analysis (INA) is a new framework for evaluating the likely success of regional interventions before, during, and after projects, for project implementers, policy makers, and funders. INA integrates across three key system components in a multilayer network analysis: (a) the quality of a management technology and the quality of research supporting it, (b) the socioeconomic networks through which managers learn about management technologies and decide whether to use them, and (c) the linked biophysical network for target species success or failure in the management landscape that results from managers’ decisions.

The specific objectives of this paper are (1) to introduce the INA framework and INA R package, (2) to illustrate identification of key nodes for smart surveillance, for networks where the likelihood of invasive species entry into the biophysical network at a given node may be based on information available to the corresponding node in the socioeconomic network, (3) to illustrate application of the INA framework for evaluating the likely degree of success of a project in intervention ecology, before, during and after an intervention, and (4) to illustrate the use of INA for evaluating adaptation strategies under global change scenarios with pulse and press stressors, introducing ‘adaptation functions’ for sustainability and resilience.

Examples of use of the INA package show one of the key outcomes of analyses: identifying when systems may be non-responsive to the system components that are readily changed through management decisions, to explore what additional adaptations may be necessary for intervention success.

The broader goal for the development of impact network analysis and the INA package is to provide a common framework that integrates across intervention ecology, to enhance opportunities for lessons learned across systems and scientific disciplines, to support the development of a community of practice, and to create a general platform for analysis of sustainability, resilience, and economic viability in intervention ecology applications.

## INTRODUCTION

Interventions in ecological systems can fail when project implementers do not understand which system components are limiting factors. The success of interventions depends on how effective the management methodologies are, whether a critical mass of decision makers adopts the necessary types of management, and the resulting efficacy of the management landscape. This is a common challenge for management intervention projects across applied ecology – including invasive and endangered species management, restoration, agricultural development, and public health programs, illustrated prominently by the COVID-19 pandemic – with opportunities for synergies in developing concepts across subdisciplines (Ostrom 2009; Chadès *et al*. 2011; Hobbs *et al*. 2011; Carvajal-Yepes *et al*. 2019; Lenzner *et al*. 2019; Hulme *et al*. 2020). Invasive species are key threats to ecological systems, while connectivity of reserves is often key to endangered species conservation (Hilty, Lidicker Jr & Merenlender 2012). Sustainable agricultural development often depends on technologies for managing the spread of pathogens and arthropod pests, and for supporting the spread of improved crop genotypes (Henry & Vollan 2014). Public health is supported by technologies for communicating about and using methods such as vaccination to slow the spread of disease (Manfredi & d’Onofrio 2013), and more broadly the One Health approach. Understanding how to optimize the benefits of research and data collection for each of these types of systems requires integration across three system components: (a) the type and quality of management technologies and the research underlying them, (b) socioeconomic networks that determine communication and influence about management technology use, such as networks of land managers or farmers, and (c) biophysical networks where decisions about use of technologies influence ecological outcomes, such as networks of pathogen invasion or networks of endangered species dispersal. Here, “impact networks” are defined as multilayer networks, composed of linked socioeconomic and biophysical networks, through which management may have a regional effect. This paper introduces a framework for scenario analysis (Garrett *et al*. 2018), “impact network analysis” (INA; Figure 1), and an R package that implements common scenarios for intervention ecology in which an impact network analysis can provide decision support for formulating project strategies.

**Figure 1.**
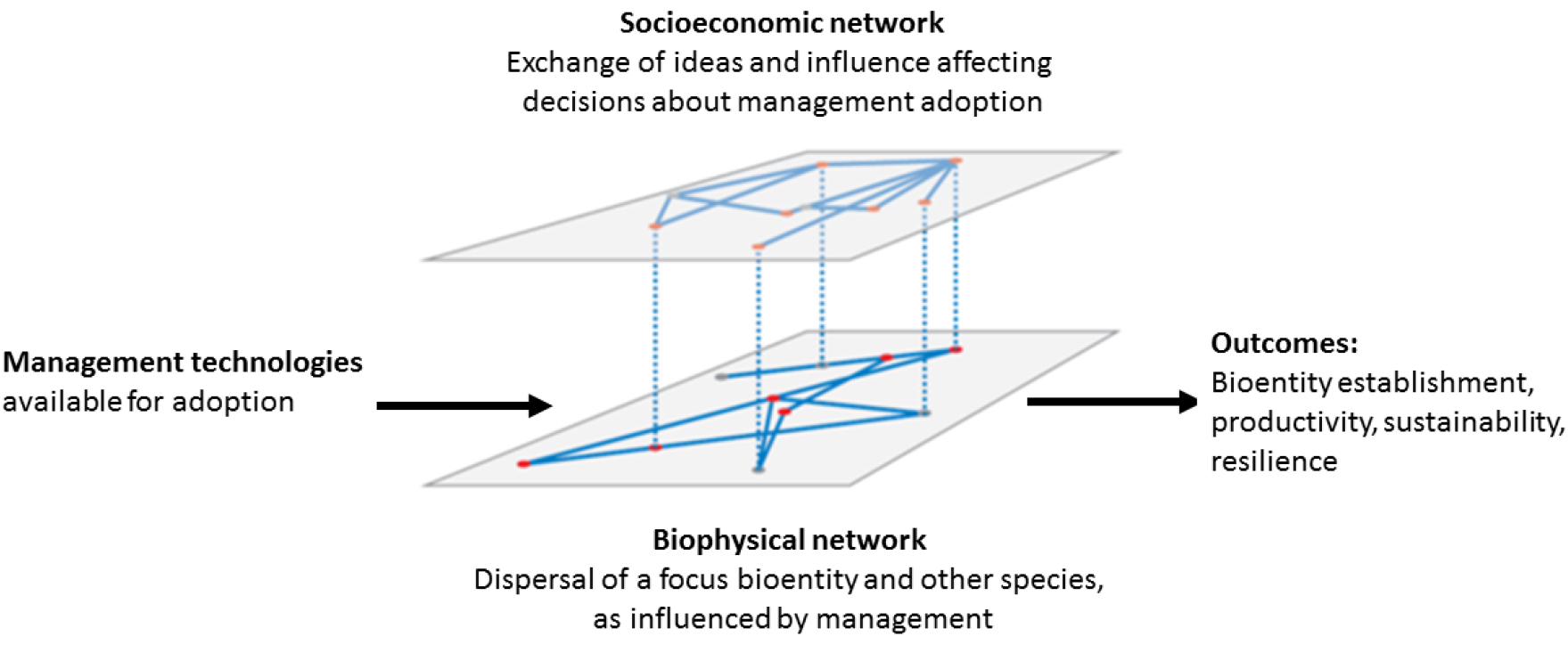
Impact network analysis is a framework for the analysis of how management technologies influence regional outcomes for management of a species or biotype (‘bioentity’) for decision support. Regional outcomes are also a function of whether decision makers are influenced to adopt management by their socioeconomic networks, and the effects of these decisions on the management landscape and its ability to effectively manage the bioentity. Nodes in the socioeconomic network are individual decision makers (such as land managers), with links in this network indicating communication and influence regarding the management technology. Some decision makers manage land nodes in the biophysical network (such as farms), indicated by a dotted line between network layers. The management technology is or is not applied at a land node, depending on the corresponding manager’s decision, and links in the biophysical network represent the potential for bioentity spread. The cumulative effects of the managers’ decisions create a bioentity management landscape, and the effectiveness of this landscape determines regional outcomes. (Figure adapted from Garrett et al. (2018) with permission from Annual Reviews.)

The first component in this framework is an intervention technology, which might be, for example, biocontrol agents, biocides, burning regimens, models indicating the best timing of management activities, or some combination of such technologies. These intervention technologies are all the products of scientific research, and can be thought of in terms of the information resulting from scientific experiments, with an associated uncertainty (Klerkx, Aarts & Leeuwis 2010). The ‘value of information’ – the improvement in outcomes when decision makers take into account information, versus not having the information – is a useful concept for regional management strategies. Analyses of the ‘value of information’ have been incorporated in, for example, medical decision making at multiple scales (Bartell *et al*. 2000; Tappenden *et al*. 2004; Claxton & Sculpher 2006), management of species (Wiles 2004; Tallis & Polasky 2011; Canessa *et al*. 2015), and adaptive resource management (Williams, Eaton & Breininger 2011). As the reproducibility of science is critically evaluated in multiple disciplines, the quality of information is a focus (Ioannidis 2005; Kenett & Shmueli 2014; Leek & Peng 2015). And even if information and technologies are of very high quality, their influence on system-level outcomes will be minimal if decision-makers are unaware of them or are not persuaded that they are a good investment of resources. Impact network analysis (INA) can be thought of as an evaluation of the realized regional value of information (including technologies broadly) in landscapes.

The second component is the socioeconomic network, where nodes are key decision makers such as farmers, other land or water resource managers, or individuals managing their families’ health (Rebaudo & Dangles 2011; Rebaudo & Dangles 2013; Burgess *et al*. 2020) – and potentially also include other agents such as scientists (Ekboir 2003), extension agents, policy makers, consumers, and related institutions. Links between nodes may indicate the spread of ideas, influence, and/or money. Individual decision-making about whether to adopt new technologies plays out in the context of the information available through individuals’ networks (Rogers 2003; Garrett 2012). Agricultural management is often limited by lack of information (Parsa *et al*. 2014), and in general heuristics for decision-making may or may not be well-developed (Ascough *et al*. 2008; Gigerenzer & Gaissmaier 2011). The effects of decision-making by agents in the socioeconomic network, with or without full information about options, creates a management landscape that influences the success or failure of species in the biophysical network.

In the biophysical network, the third component, nodes indicate the entities or geographical locations where success or failure occurs (Calabrese & Fagan 2004; Galpern, Manseau & Fall 2011). Nodes might be groups of people (as hosts to human pathogens), farms, habitat patches, or other land management units. Links between nodes indicate the potential for the spread of undesirable species or genotypes, such as antibiotic resistant human or agricultural pathogens (Margosian *et al*. 2009; Epanchin-Niell *et al*. 2010; Sutrave *et al*. 2012; Xing *et al*. 2020), or of desirable species or genotypes, such as endangered species or improved crop varieties (such as orange-fleshed sweetpotatoes to support Vitamin A consumption, e.g., evaluated in Andersen et al. (2019)). In some cases, the same type of biophysical network model may usefully be applied to related abiotic processes, such as the spread of pollutants, soil erosion, and provisioning of fresh water (Baron *et al*. 2002). Nodes in the biophysical network are linked to the corresponding decision-makers in the socioeconomic network layer, such that the probability of successful management at a biophysical node is modified by the corresponding decisions about management. Successful management also depends on the quality of information and other technologies that may be applied at a given biophysical node.

Combining these three components provides a systems perspective for decision support in scenario analyses to evaluate potential outcomes from research investments – before, during, or after projects begin. INA can also be used to evaluate the likely degree of success of adaptation strategies to pulse (intermittent) or press (continual) system stressors, such as the introduction of a new pathogen or climate change (Harris *et al*. 2018), to evaluate system sustainability, resilience, or economic viability. Some of these system components have been considered together more-or-less explicitly in disease ecology (Funk *et al*. 2009; Harwood *et al*. 2009; Funk, Salathe & Jansen 2010; Garrett 2012; Sahneh, Chowdhury & Scoglio 2012; Manfredi & d’Onofrio 2013) and natural resource management (Epanchin-Niell & Hastings 2010; Bodin & Prell 2011; Mills *et al*. 2011; Rebaudo & Dangles 2011; Hernandez Nopsa *et al*. 2015). Combining the components in an agent-based model also provides a new perspective on the science of science policy (Fealing *et al*. 2011) by directly evaluating interactions among agents engaged in developing scientific results and in implementing the new results.

The overall goal for the development of impact network analysis is to provide a common framework that integrates across intervention ecology, to enhance opportunities for lessons learned across systems and scientific disciplines, to support the development of a community of practice, and to create a general platform for analysis of sustainability, resilience, and economic viability in these types of intervention ecology applications. Integrating network analyses, as compared to more aggregated models, allows consideration of the role of geographic and social structures in the likelihood of success of technological innovations. INA is designed to provide decision support to implementers, funders, and policy makers about the prioritizations they must consider, as a complement to traditional approaches to monitoring and evaluation.

The specific objectives of this paper are (1) to introduce the INA framework and INA R package, (2) to illustrate identification of key nodes for smart surveillance, for networks where the likelihood of invasive species entry into the biophysical network at a given node may be based on information available to the corresponding node in the socioeconomic network, (3) to illustrate application of the INA framework for evaluating the likely degree of success of a project in intervention ecology, before, during and after an intervention, and (4) to illustrate the use of INA for evaluating adaptation strategies under global change scenarios with pulse and press stressors, introducing adaptation functions for sustainability and resilience. These experiments illustrate how INA can be used for analysis of hypothetical systems, observed systems, or a blend of hypothetical and observed. Data limitations will always be a challenge for scenario analyses, but uncertainty quantification methods, as illustrated here, can inform decisions about investments in interventions. Because of the complexity of most ecological systems, scenario analysis platforms like INA are needed for evaluating the likelihood of intervention success before, during, and after implementation.

## METHODS

Many applications of impact network analysis would include a combination of observed data along with simulated data that (a) represents scenarios being considered or (b) is part of an uncertainty quantification for parameters that are difficult to estimate. Three simulation experiments are presented here to illustrate use of impact network analysis for both purposes. The first is simpler, drawing on information about a biophysical network structure describing potential spread of an invasive species, where the socioeconomic network structure is implicit through weighting options for where the invasive species is likely to enter the biophysical network. This illustrates use of the smartsurv function in the INA R package. The other two experiments use the INAscene function and illustrate scenario analyses for linked socioeconomic and biophysical networks in an agent-based model of regional management of the spread of a ‘bioentity’ (a desirable bioentity, such as an endangered species or improved crop cultivar, or an undesirable bioentity, such as pathogens and other invasive species). More details about using the INA R package are available in the user guide (S1).

### 2.1 Experiment 1. Identifying key sampling locations for smart surveillance

Surveillance strategies can be informed by knowledge about the structure of the biophysical network of invasive spread. In the case of disease risk in networks for crop seed dispersal, nodes that function as hubs (high node degree) and bridges (high betweenness) in the biophysical network will tend to be important for sampling (Andersen *et al*. 2019), and as networks become more complex other node and network traits may become important (Holme 2017; Holme 2018). The relative risk of nodes being the first point of introduction of a pathogen in a network may also be a function of their role in communication networks, and communication status may be useful to include as a risk factor even when the structure of the communication network is not fully understood (Buddenhagen *et al*. 2017).

The smartsurv function in the INA package in R can be used to evaluate the importance of each node for sampling to detect spread of an invasive (Table 1). This function evaluates the invasion network to find, for each node considered as a sampling node in turn, how many other nodes remain free from the invasive by the time it is detected at the sampling node. The more nodes that remain uninvaded at the time of detection, the more effective the sampling node is for identifying invasive spread while there is still time to to manage it. Sample nodes are evaluated considering each node as a potential introduction node. The smartsurv.weight function uses the output from smartsurv to evaluate the value of sampling at each node if the probability that the invasive is introduced the network can vary from one potential introduction node to another. In practice, users of the smartsurv function would often want to provide their own estimate of the network structure for their system. In this illustration, the differences among three commonly studied types of networks are evaluated. Key nodes for sampling are identified for a set of biophysical network types, with details shown in a vignette (S2). The importance of nodes for sampling is evaluated in Experiment 1 for nine simple scenarios, representing each combination of three types of networks and three types of weighting. The three types of networks are random (Erdős & Rényi 1960), small world (Watts & Strogatz 1998), and scale-free (Barabasi & Albert 1999). The three types of weighting are unweighted, weights proportional to node degree, and weights inversely proportional to node degree.

**Table 1.** Questions in simulation experiments evaluating locations to prioritize for sampling in a smart surveillance strategy, using INA function smartsurv.

| Experiment | Component | Question |
| --- | --- | --- |
| 1A | Explicit biophysical network | What locations selected for sampling are likely to allow detection of an invasive bioentity before many other locations are invaded? |
| 1B | Added implicit socioeconomic network | How is the selection of sampling locations likely to change if the probability the invasive bioentity enters the region at each location is determined by socioeconomic traits of locations? |

### 2.2 Experiment 2. Evaluating the likelihood of management success in a region, including uncertainty quantification

In Experiments 2 and 3, the INA package function INAscene was used to perform scenario analyses in simulations in the R programming environment, using the igraph package (Csárdi & Nepusz 2006) for generating network figures. Details about the agent-based model used in INAscene are in S3 and a vignette showing how the component functions of INAscene work is in S4.

First, consider a luxurious case for scenario analysis where there is a lot of high-confidence information available about the system in which a management technology is being promoted for bioentity management, illustrated in a vignette (S5). The impact network analysis evaluates the outcomes by which success of an intervention project will be judged, such as share of region invaded, health, productivity, and profit. In experiment 2A (Table 2), the analysis is performed at the planning stage of a project, to determine the probability distribution of outcomes from the project.

**Table 2.**
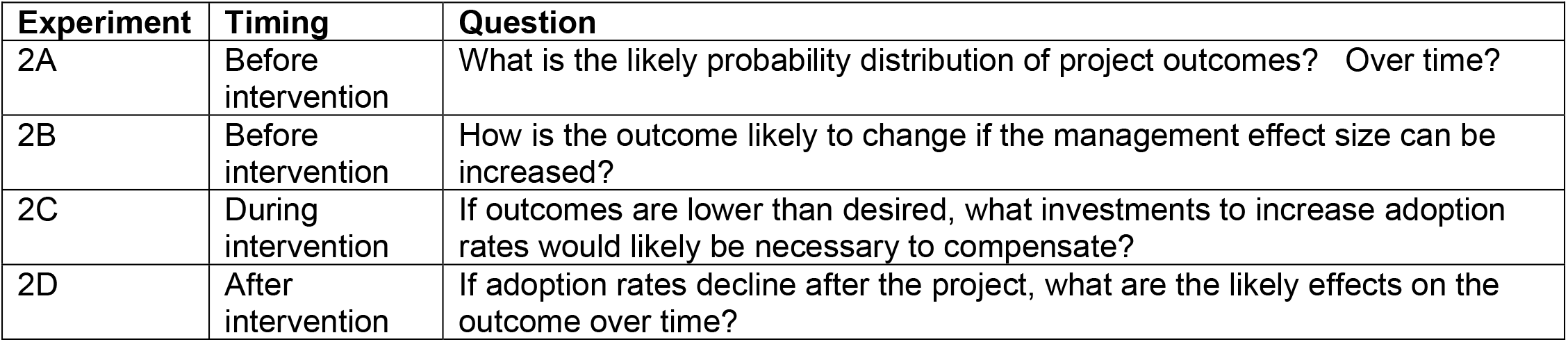
Example types of simulation experiments using impact network analysis (INA) to evaluate the likely outcomes of general **intervention projects** for managing a bioentity, using the function INAscene. The outcomes might be defined in terms of factors such as spread of an invasive species, health indicators, agricultural productivity, or the success of agricultural livelihoods.

In experiment 2B, project planners ask how the results are likely to change if the management effect size can be increased, so outcomes are evaluated for the range of possible values of the mean management effect size. This analysis is also illustrated for cases where the socioeconomic networks are based on the same common network types as in experiment 1, with the addition of a ring network for contrast.

In experiment 2C, during monitoring and evaluation of the project, suppose that the observations are consistent with the initial conceptualization of the project, but the project is performing at the low end of the initial frequency distribution of likely outcomes. If efforts are increased to enhance the probability of technology adoption, perhaps through subsidies or policies to increase uptake, what increase in adoption rates would be necessary to keep progress in the system on track?

In experiment 2D, at the conclusion of the project, the success of the project is evaluated in terms of the current status of the bioentity, and also evaluated in terms of how long the benefits of the project last for successful regional management. If adoption rates decline without project inputs (such as subsidies or educational campaigns), what happens over time? Can management effects make up for reductions in management adoption? In this example, the mean management effect size and the mean probability of adoption are varied together across their potential ranges.

#### Uncertainty quantification

Suppose there is less information available about a system. Uncertainty quantification can clarify the level of confidence in outcomes. For this case study, analyses show how system outcomes vary as a key parameter varies. Uncertainty quantification and evaluation of the outcomes for changes to the system may be evaluated similarly, with uncertainty quantification most relevant to components of the system for which it is difficult to collect data. For this case study, a parameter is varied in the inverse power law model used to describe the likelihood of movement as a function of distance.

### 2.3 Experiment 3. Adaptation to global change scenarios, including a science of science perspective

This experiment illustrates an analysis of how to modify system components that are potentially under managers’ control to compensate for changes outside managers’ control, such as changes in the likelihood of establishment – due to climate change or changes in the functional traits of the bioentity being considered. In this analysis, climate change effects are represented by changes in environmental conduciveness to establishment of a bioentity, reflected in the probability of establishment. Details of these analyses are in a vignette (S6)

In experiment 3A (Table 3), the ‘adaptation for sustainability’ scenario, conduciveness to establishment increases and remains steady over time, as a press stressor. The probability of establishment at a baseline of 0.5 is compared to a new scenario with the probability of establishment at 0.9, where this is the probability of establishment without management.

**Table 3.** Example types of simulation experiments using impact network analysis (INA) to evaluate the likely outcomes of strategies for **adaptation to global change effects** on bioentities, using the function INAscene. The outcomes might be defined in terms of factors such as spread of an invasive species, health indicators, agricultural productivity, or success of agricultural livelihoods.

| Experiment | Type of stressor | Question |
| --- | --- | --- |
| 3A | Press | To adapt for sustainability in the face of increased establishment risks, what level of change is needed in manageable system components? |
| 3B | Press | What is the **adaptation function for sustainability** describing the needed system change to keep the system failure rate below a threshold, as establishment risks increase? |
| 3C | Pulse | To adapt for resilience in the face of a rapid change in bioentity establishment, what level of change is needed in manageable system components? |
| 3D | Pulse | What is the **adaptation function for resilience** describing the needed system change to keep the failure rate below a threshold, if there is a rapid change in establishment? |

In experiment 3B, we consider an ‘adaptation function for sustainability’, the required change in a manageable component of the system to maintain system function when an unmanageable component of the system is changing. For the scenario of adaptation to higher establishment rates in the absence of management (such as due to environmental changes or changes in the functional traits of the bioentity), we consider the change in the mean adoption probability and the mean management effect necessary to compensate and keep the rate of ‘observed’ establishment at the same level as before. In this scenario, the mean probability of establishment in the absence of management for an invasive increases from a baseline of 0.5 and the goal is to keep the ‘observed’ establishment proportion below 0.2 for sustainable management, i.e., at an establishment rate no higher than under baseline conditions.

In experiment 3C, an ‘adaptation for resilience’ scenario with a pulse stressor, a baseline starting proportion of nodes infected of 0.05 is compared to a new scenario where the starting proportion has leapt to 0.50. What increase in the mean probability of technology adoption would be needed to bring the mean ‘observed’ establishment back down to the level for the baseline before the leap occurred?

In experiment 3D, the ‘adaptation function for resilience’ indicates the adaptation required in terms of modifying the system parameters under managers’ direct control to bring the ‘observed’ establishment rate back to the baseline level before the pulse stressor. This pulse stressor results in an unusually high proportion of locations with an invasive bioentity, and the system must then compensate if it is to be resilient. What adaptation is necessary to bring the proportion locations with the bioentity established below 0.2 during the time steps considered? Suppose the management effect mean is brought up to 0.9. What is the adaptation function for resilience, based on adaptation through modifying the mean technology adoption probability?

#### Science of science experiment

In a science of science scenario analysis, the ability of a technology to improve regional management is also influenced by the outcome of an initial experiment. This could represent a scenario where a research group is testing management technologies and deciding whether to promote them or not. Depending on the effort invested by the scientists in the scenario, the management effect size is estimated with greater or lesser precision. When the management effect size estimate generated by the research group is below a threshold, information about the management is not communicated, so some share of scenario realizations does not include use of the management technology. In this case study, the threshold for communication about management ranges from 0 (communication occurs regardless of the estimated management effect) to 1 (communication cannot occur unless there is not uncertainty about the complete effectiveness of management). In the first scenario analyzed here, the management effect mean is 0.5 and the management effect standard deviation is fairly high, also 0.5, while the sampling effort is low (1). Additional examples are in S6. Note that these analyses explore the potential costs of <u>not</u> communicating about a technology. There are other types of costs of communicating about a technology for which the benefits have been overestimated, where both underestimation and overestimation of technology effects may be part of a reproducibility problem in science.

## RESULTS

These results illustrate some of the types of analyses that can be implemented with the INA package, where Experiment 1 used the smartsurv function and Experiments 2 and 3 used the INAscene function.

### 3.1 Experiment 1. Identifying key sampling locations for smart surveillance

For the simple *random network* example, nodes more important for sampling for detection occur in several parts of the network, but not in the periphery (Figure 2).When weighting is proportional or inversely proportional to node degree, the relative importance of nodes shifts, as illustrated in a vignette (S3). For the *small world network* example, nodes are of similar importance, though some nodes that link across different parts of the network are somewhat more important (Figure 2). When weighting is proportional or inversely proportional to node degree, there is little change because of the similar roles of nodes (S2). For the simple *scale-free network* example, the high degree nodes are clearly more important for sampling (Figure 2). When weighting is proportional or inversely proportional to node degree, there is again only a slight change because the role of high degree nodes in driving the invasion network is so important (S2). When stochastic networks are considered, the clear importance of some nodes for sampling in deterministic networks is decreased (S2).

**Figure 2.**
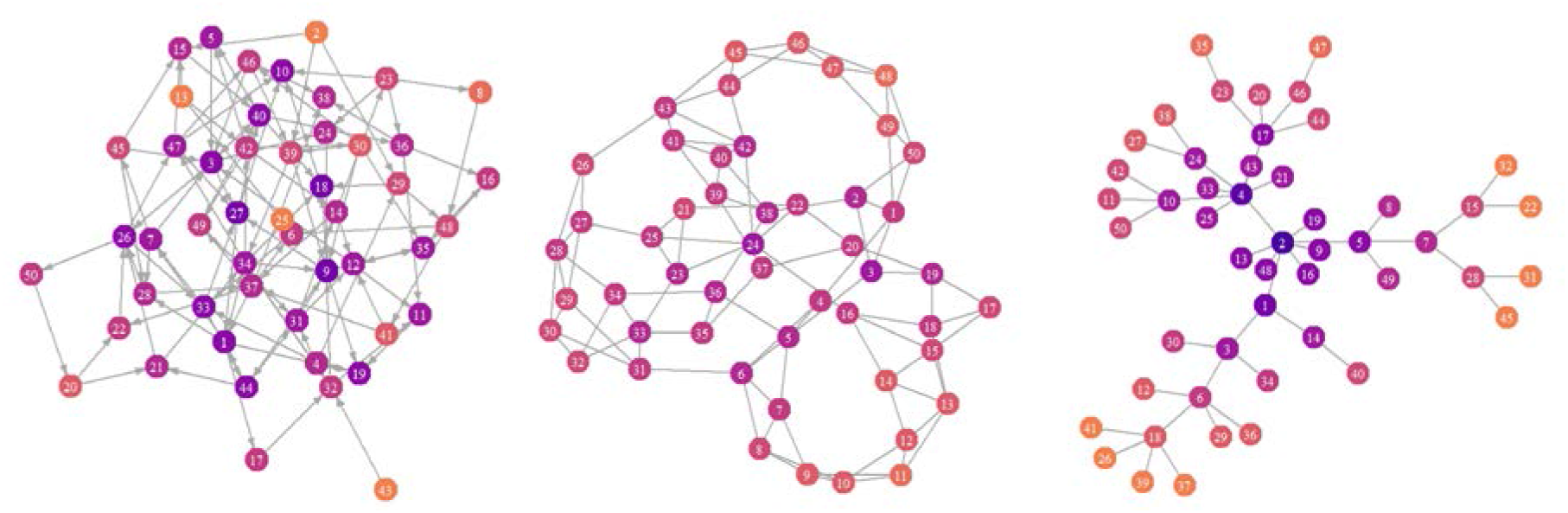
Identifying key nodes for sampling for an invasive ‘bioentity’ as part of a smart surveillance strategy (Experiment 1). This analysis identifies the nodes where sampling can detect the bioentity before it has spread further through the network. In these simple examples based on commonly-studied networks, darker nodes would detect the epidemic sooner, while detection at lighter nodes would occur when there were fewer opportunities for managing the epidemic. (Left) In a **random network**, nodes at the periphery of the network will be less important for sampling. (Center) In a **small world network**, nodes may be of similar value for sampling, but those that link across sections of the network may be more useful. (Right) In a **scale-free network**, nodes with high degree are more useful for sampling. As real networks diverge from these simpler structures, and become too large for simple visualization, this method for identifying key nodes for sampling can be used to find important nodes.

### 3.2 Experiment 2. Evaluating the likelihood of management success in a region, including uncertainty quantification

In Experiment 2A, at the planning stage of a project, analysis of the probability distribution of outcomes from the project indicates that the bioentity would become established in less than half the nodes (Figure 3, with more details in S5).

**Figure 3.**
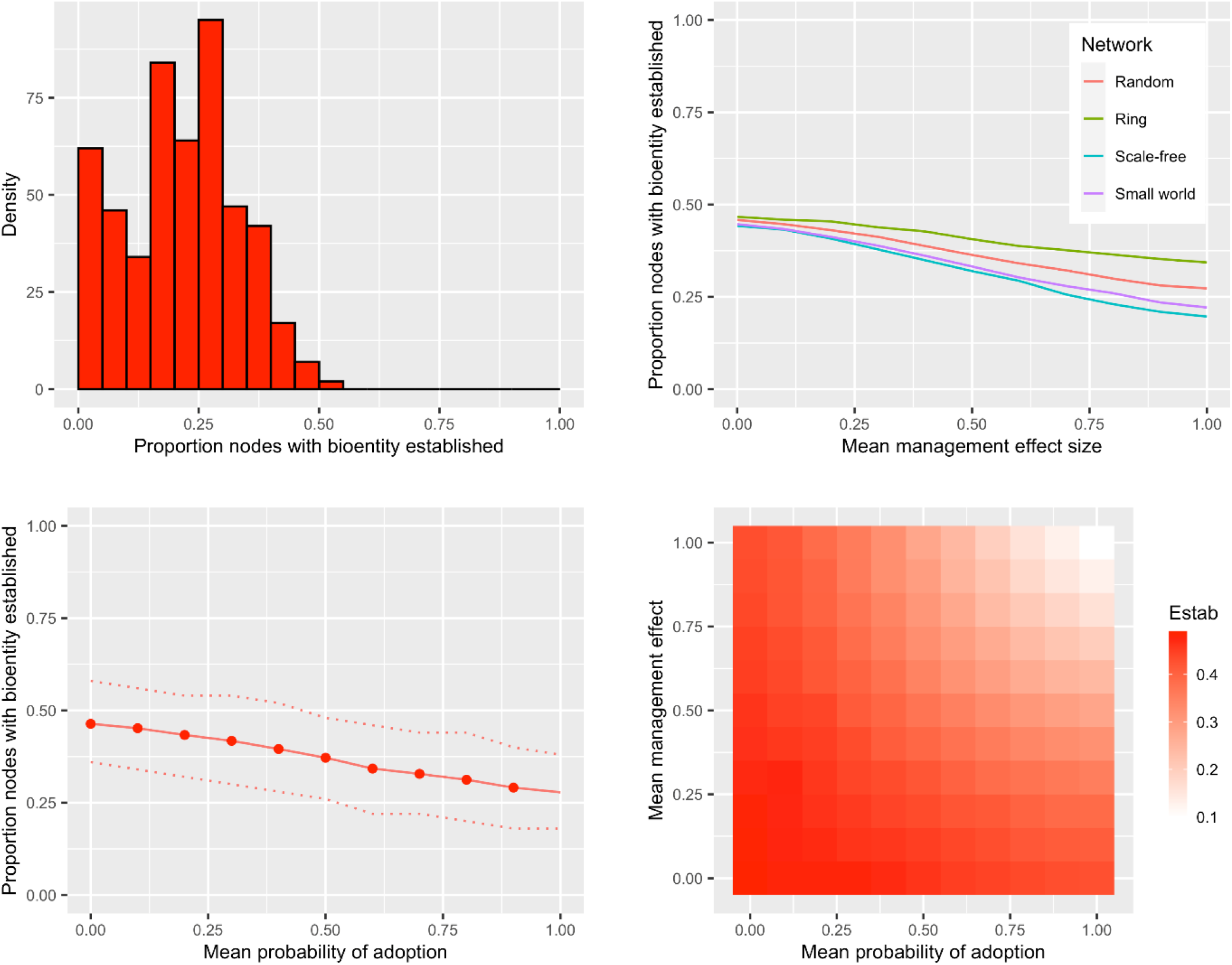
Evaluating the likelihood that management is successful in a set of scenario analyses (Experiment 2, with more details in S5) (Upper left) At the outset of a project, the analysis focuses on the likely distribution of the ‘observed’ proportion of nodes with the ‘bioentity’ established. (Upper right) If the project must be adjusted when performance needs to be improved, the analysis focuses on the response of the system to changes in the mean management effect size, illustrated here for four different types of socioeconomic networks in what is otherwise the same system. (Lower left) If the performance of the system is weak, the analysis focuses on whether the results can be improved by boosting performance in a feature such as the mean probability of adoption (where dotted lines indicate the 5^th^ and 95^th^ percentile of simulation results). (Lower right) In consideration of how project benefits can persist over time, the analysis focuses on what combinations of ‘manageable’ components are needed – in this case, the mean management effect size and the mean adoption rate – to keep the ‘observed’ establishment rate acceptably low for an unwanted bioentity (e.g., the dark red range).

In Experiment 2B, network types are compared in terms of how responsive the system is to changes in the management effect size. The small world network generally has a greater benefit from increasing the management effect, while the ring network generally has the least benefit of the systems considered (Figure 3).

In Experiment 2C, the system has a weak response to changing the mean probability of adoption (Figure 3), so that even when adoption is certain (for nodes with information about the management technology), other changes in the system would be necessary to keep the ‘observed’ establishment rate below 0.2.

In Experiment 2D, evaluating how benefits of a project intervention can persist over time, a combination of higher mean management effect size and higher mean adoption rate can push the ‘observed’ establishment rate below 0.2 (Figure 3).

In an uncertainty quantification, the effect of changing a power law parameter (determining the dispersal gradient for the bioentity) was evaluated, representing the types of parameters that may often be difficult to estimate. For the scenario considered, the ‘observed’ rate of establishment is similar across ranges of the parameter between 0 and 0.7 and between 1.2 and 2.0 (Figure 4). However, if the true parameter value is between 0.7 and 1.2, obtaining a more precise estimate of the parameter may be important for understanding system outcomes.

**Figure 4.**
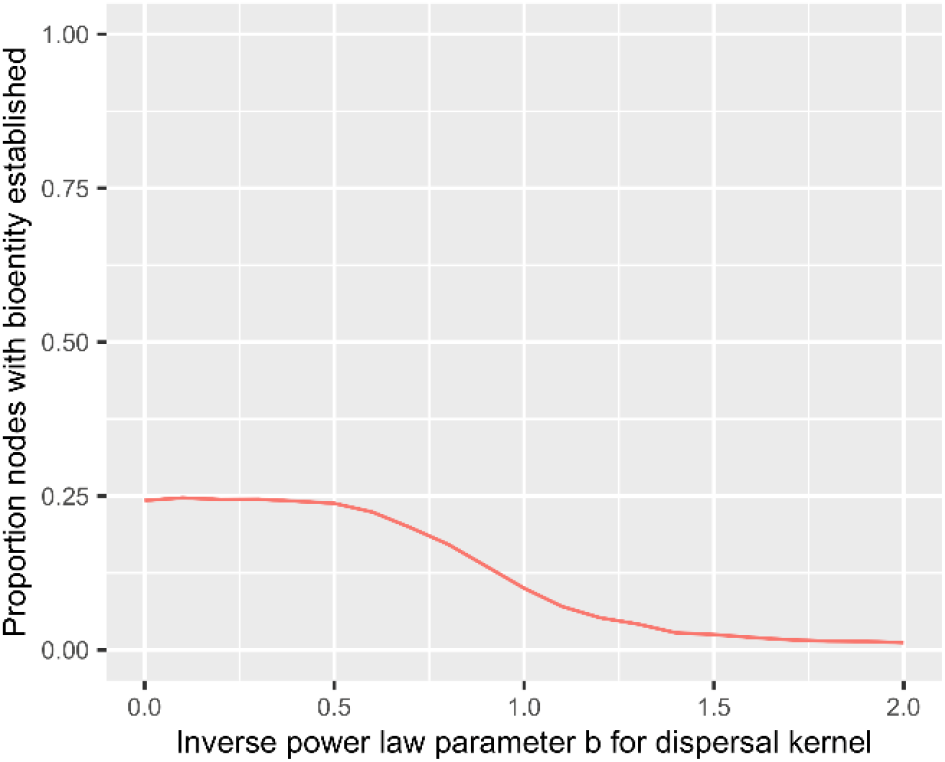
An uncertainty quantification to evaluate the effect of changing am unknown power law parameter describing the dispersal gradient for a bioentity.

### 3.3 Experiment 3. Adaptation to global change scenarios, including a science of science perspective

In Experiment 3A, the probability of establishment increases from a baseline of 0.5 to a new scenario of 0.9, and both the probability of adoption and the management effect size must change to compensate (Figure 5, with more details in S6).

**Figure 5.**
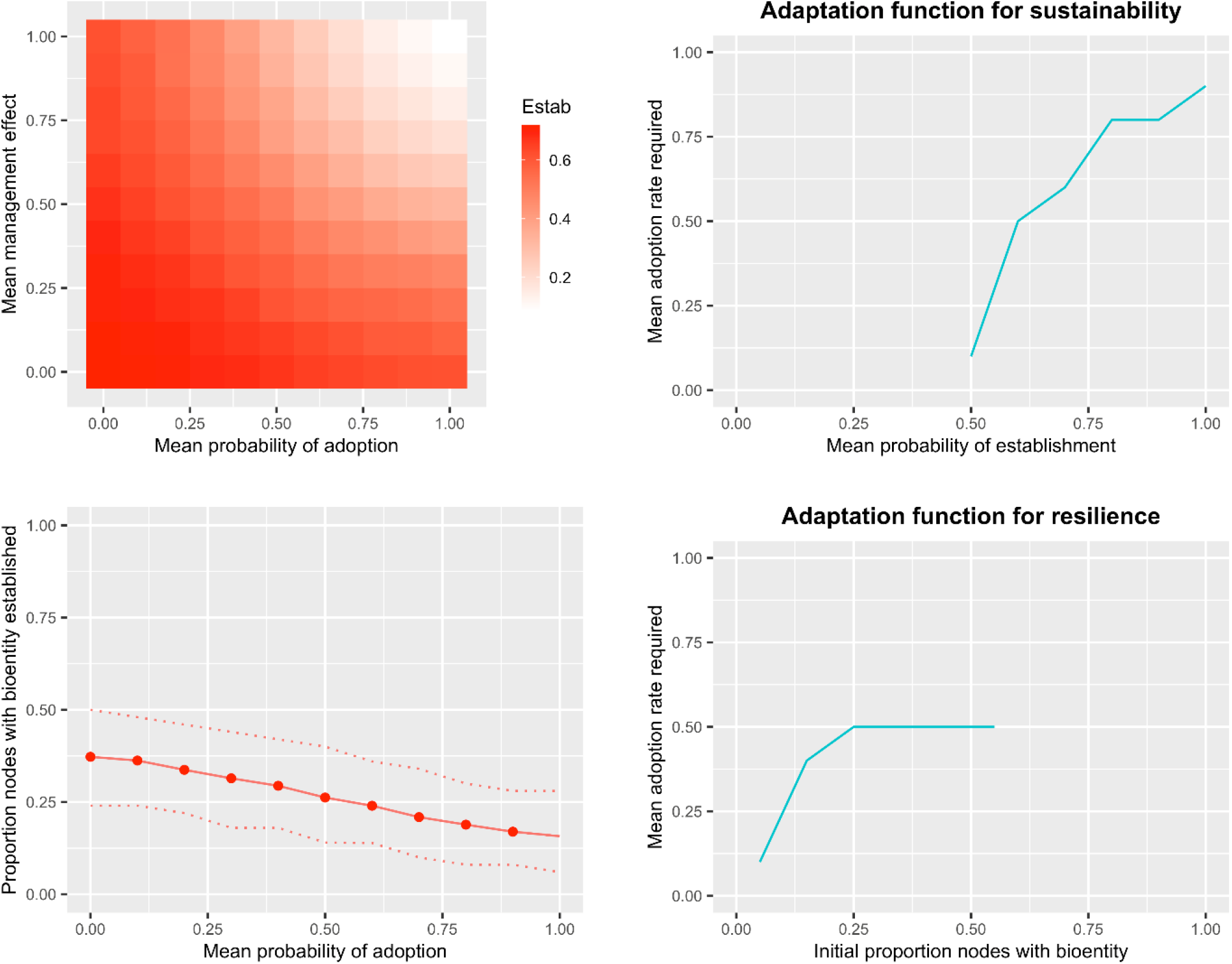
Sustainable or resilient adaptation requirements in global change scenarios (Experiment 3). (Top left) The sustainability of a system – in terms of its ability to keep bioentity establishment (‘Estab’) below a threshold even as a press stressor increases the probability of establishment – is evaluated as the probability of establishment moves from a baseline of 0.5 to a new level of 0.9 and both the mean management effect size and the mean probability of adoption increase to compensate. (Top right) The ‘**adaptation function for sustainability**’ indicates the required change in a manageable component of the system (mean probability of adoption of a management technology) to keep the ‘observed’ establishment rate at the baseline level as the mean probability of establishment (in the absence of management) increases. (Bottom left) The resilience of a system – in terms of its ability to return bioentity establishment to a baseline level after a pulse stressor elevates establishment – is evaluated in terms of how well adjusting the mean probability of adoption can compensate. (Bottom right) The ‘**adaptation function for resilience**’ indicates the required change in a manageable component of the system (mean probability of adoption of a management technology) to return the ‘observed’ establishment rate to the baseline level after a pulse stressor boosts the establishment.

In Experiment 3B, the adaptation function for sustainability is evaluated. This function indicates the required change in a manageable component of the system to maintain system function when an unmanageable component of the system changes. The adaptation function for sustainability shows how the mean adoption rate must change (given also an increase in the management effect size) to compensate for changes in the mean probability of establishment (Figure 5, S6).

In Experiment 3C, adaptation for resilience is evaluated in response to a pulse stressor that pushes the baseline proportion of nodes with the bioentity from 0.05 to 0.50. Increasing the mean probability of adoption can keep the ‘observed’ establishment proportion down about to 0.2 (Figure 5), but more adaptation would likely be necessary to keep the establishment proportion reliability lower.

In Experiment 3D, the adaptation function for resilience is evaluated. This system recovers from the pulse stress of 0.50 nodes with the bioentity. The adaptation function for resilience when responding to the pulse stressor evaluates the technology adoption rates required to compensate for an increasing initial number of nodes with the bioentity established (Figure 5 and S6).

In science of science experiments, as the management effect size threshold for communication about the management technology increases, the ‘observed’ rate of establishment increases, as expected (S6). For the second case in which the probability of establishment and the probability of adoption are both 0.9, when the threshold is between 0.2 and 0.8, the uncertainty in the outcome is high, as the simulated experiment to evaluate the technology may or may not be adequate to ascertain that the technology is worthy of communication (Figure 6).

**Figure 6.**
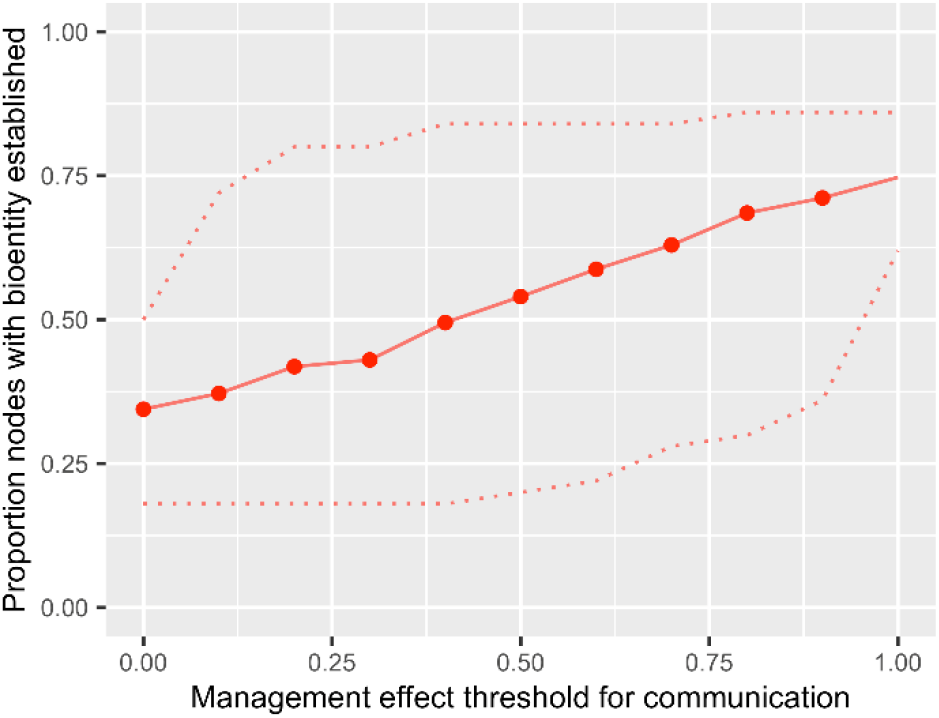
A science of science experiment to evaluate the effects of changes in a communication threshold, where a simulated experiment may result in estimates of the management effect size above or below the threshold, as a function of the true management effect size, the variance in the effect, and the research effort invested. Dotted lines indicate the 5^th^ and 95^th^ percentile of simulation results.

## DISCUSSION

These case studies illustrate some of the potential applications of INA, in general, and how to use the INA R package. In the examples, all components such as initial distributions and networks are simulated following ideas about their structures in hypothetical scenarios, while many INA applications would use a mixture of observed and simulated data. The Discussion presents ideas for expanding on these types of experiments with observed data and a mixture of scenario analyses and uncertainty quantification to address missing information. The INA R package is intended to expand in future versions to incorporate new types of related analyses, and some ideas follow about useful future applications. Even the simple scenarios above illustrate a couple general points that these types of analyses can reveal. One point is the potential for thresholds in response to stressors, a problem often discussed. Another point is the potential for flat responses to adaptation strategies, which is less commonly considered but can be an important challenge for wicked problems in regional management. Projects may stall until the components of the system that are limiting factors are understood, as a system may be insensitive to changes in readily managed components. Use of INA to identify limiting factors and necessary adaptations can be expanded by integrating relevant data layers, such as maps of environmental conduciveness or satellite images with information about the locations of species, in a more detailed decision support system. There is often a critical window of opportunity for management in impact networks, and efficient INA applications have the potential to generated automated updates about likely outcomes from the regional management strategies being considered.

In analyses of the potential roles of nodes in a smart surveillance strategy (using the function smartsurv from the INA package) in Experiment 1, the results for the hypothetical cases illustrate patterns of node importance for surveillance in typical random, small world, and scale-free networks (Figure 2). In analyses of the value of locations for surveillance, the socioeconomic network may only be implicit, in terms of likely entry points into the biophysical network, as in Buddenhagen et al. (2017) where nodes with less reliable information sources may be more likely pathogen entry points. Other reasons for higher risk of being the starting node might result from a node’s role as a port, weather conditions associated with a node, or lack of resources for management at the node. In new applications of smart surveillance analyses, specific networks representing systems may have unique properties (Holme 2017; Holme 2018). Users can evaluate the importance of nodes in their specific biophysical networks, and potentially include new analyses such as evaluating whether node demographic or other traits are associated with higher or lower importance for surveillance. Analyses such as evaluating the connectivity of host populations (Xing *et al*. 2020), can be used to characterize landscapes for application with smartsurv. In on-going surveillance analyses, the importance of nodes could be updated as more information about the system becomes available.

Analyses of how well an intervention project is likely to succeed at regional management of a bioentity (using the function INAscene from the INA package) are illustrated in Experiment 2. The results for hypothetical cases show how management options singly or in combination may be adjusted to make projects more likely to succeed, and how network structures may modify system responsiveness to management effect size (Figure 3). Changes in the management effect size might be attainable through further experimentation, and the probability of technology adoption might be modified through policies such as subsidies. In applications using INAscene, users could provide a combination of observed and hypothetical components to evaluate the likelihood of project success, updating as new information becomes available. Analyses might evaluate the likely outcomes for categories of nodes, perhaps testing hypotheses about how well systems perform as a function of manager traits such as gender or wealth, or as a function of node traits such as environmental factors. Similarly, scenario analyses might evaluate potential new impact network structures and how these structures benefit different types of nodes. The results of analyses can help to inform project corrections. The illustration of uncertainty quantification shows how the effect of parameters that may be challenging to estimate with precision, such as parameters describing dispersal gradients, can be evaluated (Figure 4). It is convenient when the results of scenario analyses are similar across parameter values in uncertainty quantification (e.g., in Andersen et al. (2019)), but if not, it is still helpful to know what new data would be particularly valuable to collect for understanding the system better.

Achieving sustainability and resilience of systems and their ecosystem services is a key challenge, and often operationalizing these concepts is an additional challenge for interventions to try to achieve these goals (Howden *et al*. 2007; Clark *et al*. 2011; Biggs *et al*. 2012; Standish *et al*. 2014). In analyses of potential adaptation strategies to maintain sustainability or resilience, with regard to regional management of a bioentity (using the function INAscene from the INA package) in Experiment 3, the results for hypothetical cases introduce evaluation of adaptation functions. Adaptation functions are defined here to represent the change in one (or more) system parameters necessary to achieve desired system outcomes. Adaptation functions for sustainability indicate the changes in manageable parameters necessary to return a system to the outcomes before a press stressor; adaptation functions for resilience indicate the changes in manageable parameters necessary to return a system to the outcomes before a pulse stressor (Figure 5). For new use of the INAscene function to study adaptation strategies, a combination of observed and hypothetical data could be input. Uncertainty quantification might include uncertainty about the magnitude of global change factors, with an emphasis on minimizing the lag time in response to stressors. Other types of global change, such as increased trade, might lead to other types of system change, such as biophysical networks with more links. The science of science experiment illustrates the effects of decision-making about communication based on research results (Figure 6). Variations in the science of science experiments can address issues in the reproducibility of research, and can be integrated with ideas about the ‘theory of applied statistics’ and how to optimize statistical design for regional management benefits.

The priorities for defining scenarios will differ from one study to another. When evaluating the likely success of interventions that are under immediate consideration, analyses will often try to achieve the greatest level of precision possible given the data available. When considering the potential for specific types of future interventions, or the theory of effective interventions, other priorities may be at least as important. There are often trade-offs in the ability of a model to achieve precision, realism, and generality (Levins 1966; Gross 2013). Other applications of impact network analysis could focus on developing general theories for the development of future intervention strategies (Figure 7). Agroecological seed systems are an important example of multilayer networks supporting agricultural sustainability and resilience. Layers include the network of seed movement in formal and informal systems, the network of pathogen or pest movement through seed, and the network of information and influence related to integrated seed health strategies (Thomas-Sharma *et al*. 2016; Thomas-Sharma *et al*. 2017). Successful seed systems will optimize the maintenance and spread of desirable crop varieties (Pautasso *et al*. 2013; Pautasso 2015; Labeyrie *et al*. 2016) while minimizing the spread of pathogens through seed or grain movement (Hernandez Nopsa *et al*. 2015; Buddenhagen *et al*. 2017; Andersen *et al*. 2019). Additional linked networks include the global network of crop breeders who exchange genetic material (Garrett *et al*. 2017). Hypothetical networks could be generated using methods such as exponential random graph models (ERGMs) to test hypotheses about system outcomes for different ERGMs (Lusher, Koskinen & Robins 2013). Another type of research could focus on the effects of combining different commonly studied network models for socioeconomic networks and biophysical networks. For example, a socioeconomic network might start with a distancebased probability of link existence as for the corresponding biophysical network, but then have a given probability of re-wiring. Theories about the likelihood of regional management could be developed in terms of the relationship between the socioeconomic and biophysical networks, with applications for new regional management scenarios.

**Figure 7.**
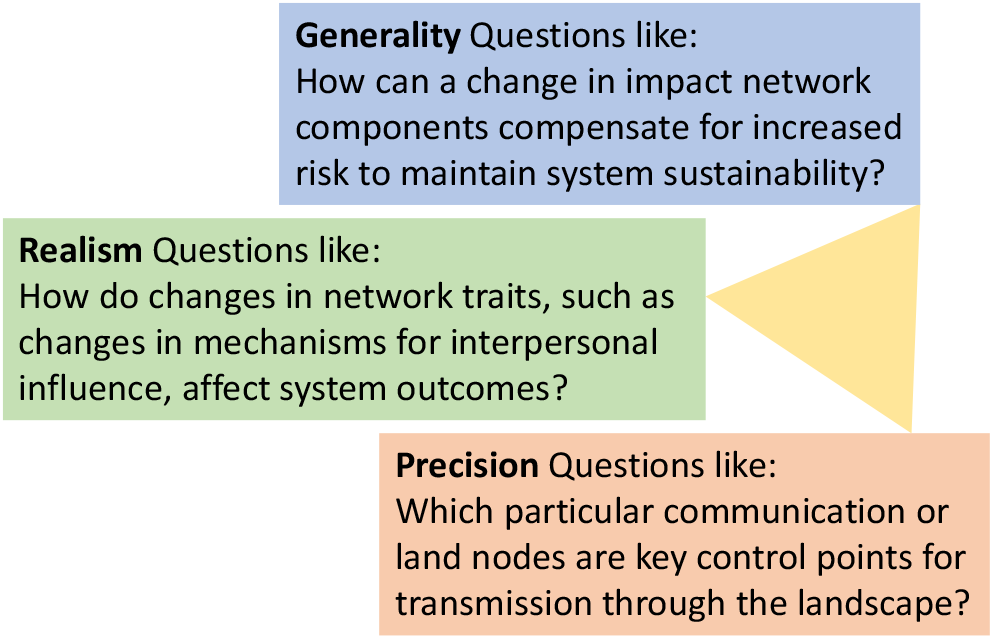
Three potential priorities in impact network analysis, and examples of the types of questions that might be asked in each context. As more information about a system is available, questions can address greater realism and precision.

The INA package is designed to be the basis for future expansions to better address specific types of systems. For smart surveillance strategies using the smartsurv function, next steps will include explicit analyses of specific types of sampling strategies and their relative performance for sets of specified starting locations for the bioentity. For scenario analyses using the INAscene function, next steps will include options for tracking population sizes at nodes. Another focus will be generation and evaluation of observations of management success in a heterogeneous landscape, addressing ‘big data’ in the form of information that is generated throughout, and potentially spread throughout, a network, such that managers may evaluate this information (Cui *et al*. 2016). For global change scenarios, new components will include the potential for temporal or spatial trends in parameters. For science of science scenarios, new components are planned to address the costs of research and the cost of management implementation, and to address technology upscaling and the formation and dissolution of links. Another useful extension will be consideration of multiple bioentities.

Operationalizing the concepts of sustainability and resilience are ongoing challenges (Standish *et al*. 2014) and INA is an option for evaluating the limits of responsiveness of a system and what is likely to be feasible within those limits for management adaptation. Major challenges remain for management of biodiversity while meeting needs for food production (Leclère *et al*. 2020), where communities addressing these problems have additional scientific commonalities in intervention ecology addressed by methods such as INA. Intervention ecology can also draw on advances in physics and the potential to integrate across many different types of socioeconomic and biophysical networks (Harwood *et al*. 2009; De Domenico *et al*. 2016). The broader goal of the INA framework is to support a community of practice through application across a wide range of system contexts and questions, providing research spill-over and cross-disciplinary lessons learned. As regional management strategies incorporate new approaches, including artificial intelligence for decision support, INA can be applied to integrate data layers rapidly to aim for effective management during critical periods when success is more likely.

## ACKNOWLEDGEMENTS

Development of the INA package was undertaken as part of, and funded by, the CGIAR Research Program on Roots, Tubers and Bananas (RTB), supported by CGIAR Trust Fund contributors, and by USDA NIFA grant 2015-51181-24257. I also appreciate support for concept development from US NSF Grant EF-0525712 as part of the joint NSF-NIH Ecology of Infectious Disease program; US NSF Grant DEB-0516046; the CGIAR Research Program on Climate Change and Food Security (CCAFS); USDA APHIS grant 11–8453–1483-CA; the USAID Feed the Future Haiti Appui à la Recherche et au Développement Agricole (AREA) project AID-OAA-A-15-00039; The Ceres Trust; NCR SARE Research and Education Grant LNC13-355; and the University of Florida. The contents are the responsibility of the author and do not necessarily reflect the views of these funders, or the United States Government. Thanks to Y. Xing, K. F. Andersen, R. A. Choudhury, and P. Garfinkel for helpful input. This work is dedicated to the memory of C. R. Garrett, B. B. Garrett, and J. B. Garrett.

